# Glucocorticoid-induced proteome and phosphoproteome changes in breast cancer cell lines

**DOI:** 10.1101/2025.11.25.690354

**Authors:** Hayoung Cho, Jesper V. Olsen

## Abstract

Glucocorticoids (GCs) are steroid hormones that bind to glucocorticoid receptor (GR) as ligands to initiate systemic anti-inflammatory effects. GCs are commonly administered alongside chemotherapy to reduce treatment-related side effects in breast cancer patients. However, GC administration has been shown to promote metastasis in breast cancer. In this study, we used quantitative mass spectrometry-based approaches to analyze proteome and phosphoproteome of three breast cancer cell lines following GC treatments. By comparing MCF7, MDA-MB-231, and MDA-MB-436 cells, we suggest that the level of GR significantly affects GC-mediated responses. Additionally, we identify noncanonical transcription factors (TFs) and kinases that are regulated by GCs in different cell lines. Together, our data present GC-induced modulations and modifications at protein level, indicating that breast cancer metastasis occurs via TF- and kinase-dependent signaling pathways. These findings highlight the need for careful consideration of GC use in breast cancer therapy and identify potential molecular targets for mitigating adverse effects.

**Graphical abstract:** 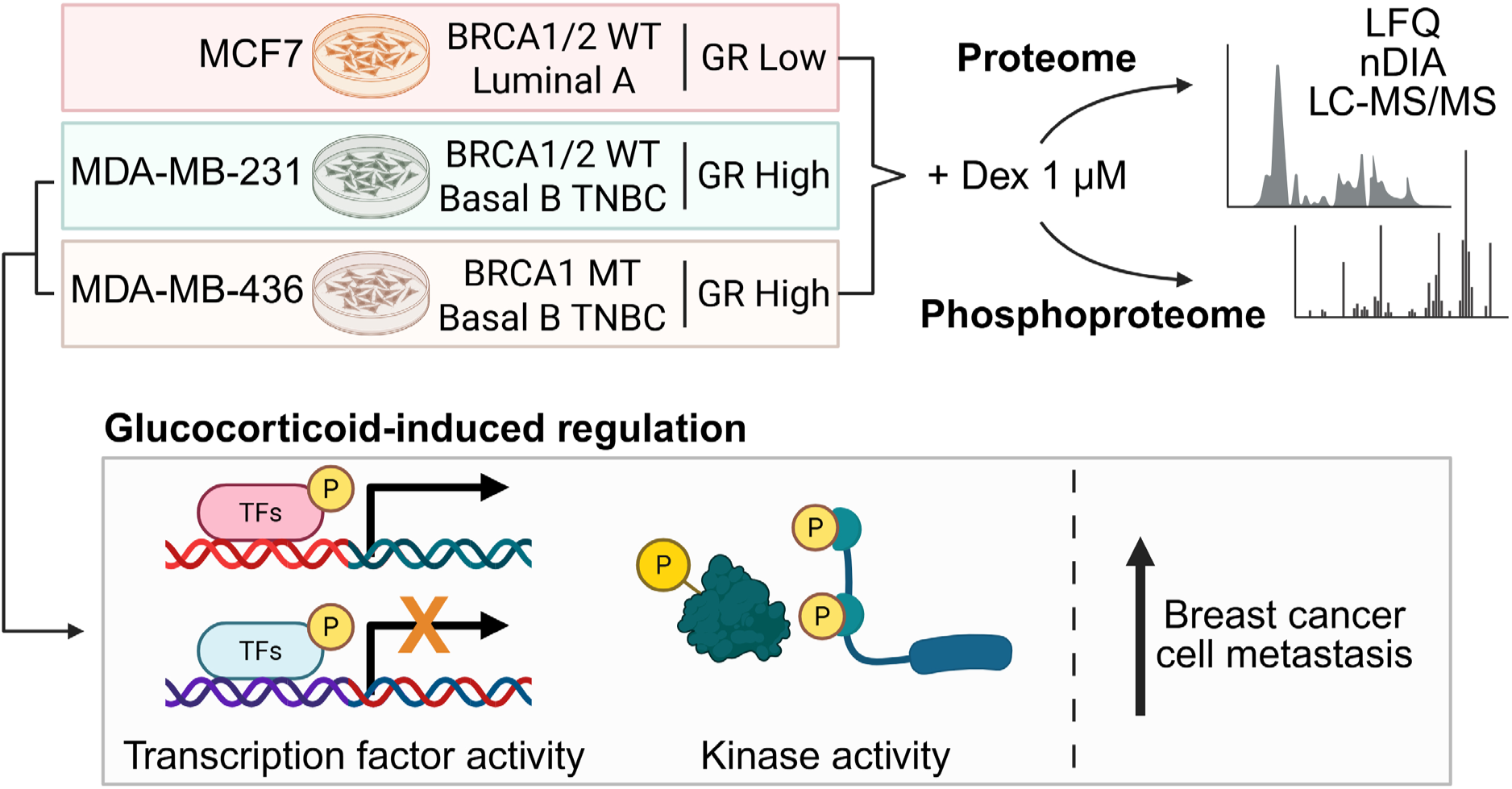

## Introduction

Breast cancer is one of the most lethal types of tumors that has the highest incidence rate in female cancer patients [1]. At molecular level, breast cancer is highly heterogenous by nature. It can be classified into four main subtypes: basal-like triple-negative breast cancer (TNBC), human epidermal growth factor receptor 2 (HER2, also known as *erbB2*)-positive (HER2^+^), luminal A, and luminal B. This is based on the expression of hormone receptors, estrogen receptor (ER) and progesterone receptor (PR), a cell proliferation marker Ki67, and HER2. Somatic mutations in genes, such as *BRCA1*, *BRCA2*, and *TP53*, also contribute to development and aggressiveness of breast cancer [2].

According to the Surveillance, Epidemiology, and End Results (SEER) program (2014-2020), 5-year relative survival rate for breast cancer patients remains high at 91% but sharply drops to 32% for those diagnosed with distant or metastatic breast cancer. Among 15 distant metastasis sites, the most commonly affected organs are bone, followed by lung, liver, and brain. By subtype, luminal tumors are more prone to bone metastasis, whereas TNBC and HER2^+^ tumors show preference for intestinal and brain metastases [3]. Data from an ongoing PRAEGNANT breast cancer registry (NCT02338167) indicate that *BRCA1* mutation is associated with more than double the rate of brain metastasis compared to wildtype tumors [4]. Similarly, a retrospective study of breast cancer patients between 2000 and 2017 found that *BRCA1* mutation increased brain metastasis in TNBC, although the overall frequency of brain metastasis was similar between carriers and noncarriers. Nonetheless, brain metastasis notably reduced survival across all breast cancer types [5]. These reports highlight the need for a deeper understanding of molecular heterogeneity and its impact on treatment responses to advance breast cancer clinical development.

Transcription factors (TFs) function as master regulators of carcinogenesis by controlling expression of many target genes and their downstream pathways. In breast cancer, TFs have pivotal roles in facilitating stem cell-like features, epithelial-to-mesenchymal transition (EMT), resistance to chemotherapy, tumor growth, and metastasis. Nuclear receptors (NRs) are one of the widely studied TFs in breast cancer, and several NR inhibitors underwent Phase I, II, and III clinical trials. Other well-established TFs, including MYC, E2F, ETS1, and β-catenin, are also known to contribute to breast cancer cell metastasis and invasion [6].

For over 40 years, glucocorticoids (GCs) have been co-administered with chemotherapy in solid tumor patients [7]. Synthetic GCs such as the FDA-approved dexamethasone (Dex) and prednisolone (Pred) help alleviate side effects of breast cancer treatments through immunosuppressive mechanisms [8]. However, the effect of GC treatment in breast cancer remains controversial. Low-dose administration of Dex suppressed tumor growth and distant metastasis in mouse models, while high-dose displayed opposite effects [9]. Additional studies support that GC treatment promotes breast cancer metastasis. GR activation by Dex or triamcinolone acetonide induced distant metastasis to spleen, ovaries, lung and liver, while enhancing tumor colonization and reducing survival *in vivo*. Integrative proteomics, phosphoproteomics, and transcriptomics further revealed that GR activation stimulated signaling pathways associated with metastasis [10]. Moreover, Dex-induced phosphorylation of GR at Ser-134 increased expression of genes involved in TNBC cell migration and metastasis, as supported by transcriptomic analysis [11]. Lastly, GC treatment mediated neutrophils to create metastasis-friendly tumor microenvironment in mouse models, promoting breast cancer dissemination to lungs [12]. Taken together, these reports raise concerns about potential pro-metastatic functions of GCs in breast cancer.

This study aims to investigate the effects of GCs in breast cancer cell lines with different molecular characteristics using quantitative mass spectrometry (MS). Global analyses of proteome and phosphoproteome changes were conducted following 1 µM Dex treatment for 24 hours in the hormone-sensitive MCF7 cell line as well as two TNBC cell lines MDA-MB-231 and MDA-MB-436. We show that the endogenous level of glucocorticoid receptor (GR) influences Dex-induced protein-level reprogramming. Additionally, Dex treatment significantly regulates the activity of several TFs and kinases in MDA-MB-231 and MDA-MB-436 cells. These alterations involve proteins associated with metastatic processes, suggesting that GC treatment may promote breast cancer cell metastasis.

## Materials and Methods

### Cell culture and treatment

MCF7, MDA-MB-231, and MDA-MB-436 cells were cultured at 37°C with 5% CO_2_ in DMEM high glucose (Gibco^TM^ 11965092) supplemented with 10% penicillin/streptomycin and fetal bovine serum (FBS). Cells were subcultured every 3-4 days and were tested for mycoplasma regularly. Cells were treated with 0.1% DMSO (Sigma-Aldrich D2650) or 1 µM Dex (Sigma-Aldrich D4902) for 24 hours before lysis.

### Mass spectrometry sample preparation

For total proteome, cells were lysed with pH 6.8 50 mM Tris-HCl buffer (100 mM NaCl, 1% Triton X-100, benzonase, protease inhibitor tablet) and sonicated at 30% amplitude for 5 seconds x2, followed by centrifugation at 16000 rpm 4°C for 15 min. 50 ug proteins were manually PAC-digested with 25 ul of hydroxyl beads (MagReSyn MR-HYX400) then stage-tipped with 4 layers of C18 disks.

For phosphoproteome, cells were washed 2x with ice-cold PBS containing phosphatase inhibitors (2 mM sodium orthovanadate, 1 mM β-glycerophosphate, 1 mM NaF). Cells were lysed with boiling 5% SDS buffer (5 mM TCEP, 10 mM CAA, 100 mM Tris pH 8.5), incubated at 95°C for 10 min shaking at 1400 rpm, and sonicated at 40% amplitude for 5 sec x2. 500 µg of proteins were PAC digested using KingFisher (Thermo Fisher Scientific) and eluted in 50 mM TEAB containing 2 µg trypsin and 1 µg lys-C. After overnight digestion, samples were acidified with 10% FA and dried by speedvac for 2 h. Samples were reconstituted in 80% ACN/5% TFA/0.1M glycolic acid (GA) and incubated with 15 µL of Ti-IMAC beads (MagResyn) and washed with 80% ACN/5%TFA/0.1M GA, 80% ACN/1% TFA, and 10% ACN/0.2% TFA using KingFisher. Phosphoenriched peptides were eluted from beads with 1% NH_3_OH, acidified with 10% TFA, and loaded onto Evotips for LC-MS/MS analysis.

### Mass spectrometry analysis

Total proteome samples were separated with Vanquish Neo UHPLC system (Thermo Fisher Scientific) coupled with Orbitrap Astral, separated with home-pulled 15 cm x 75 µM column packed with ReproSil-Pur 120 C18-AQ 1.9 mm beads (Dr. Maisch). Total proteome was separated for 30 min in narrow-window data-independent acquisition (nDIA) mode, full scan orbitrap resolution set to 240,000 in positive ion mode, ion transfer temperature (ITT) set to 275°C, RF lens 50%, normalized AGC target 250%, and isolation window (m/z) of 4. Full scan range (m/z) was set to 300-1000 with maximum injection time of 50 ms.

Phosphoproteome was analyzed on Evosep One LC system coupled with Orbitrap Astral in nDIA mode. Samples were separated with 15 cm x 150 µM performance column (EV1137; Evosep) at a gradient of 30 samples per day. Full scan orbitrap resolution was set to 240,000 in positive ion mode, ITT set to 275°C, RF lens 40%, normalized AGC target 500%, and isolation window (m/z) of 2. Full scan range (m/z) was set to 480-1080 with maximum injection time of 30 ms.

### Data processing

Total proteome and phosphoproteome MS raw files were analyzed with Spectronaut (Biognosys) v19 using human UniprotKB fasta (reviewed in May 2024) and cell contaminants in a library-free mode (direct DIA+). For fixed modification, cysteine carbamidomethylation was selected. For variable modification, protein N terminus acetylation and methionine oxidation were selected. Quantification was calculated with cross-run normalization disabled. Phosphoproteome was searched with an additional variable modification of serine, threonine, and tyrosine phosphorylation. Minimum localization probability was set to 0.75.

### Bioinformatics

Protein groups and phosphosites were filtered using Perseus v2.0.11 to keep those that had valid values of at least four per condition. Label-free quantifications were log_2_ transformed, then imputed with width 0.15 and down shift 2 in total matrix mode. Data was median normalized using R (v4.4.1) and RStudio (v2024.04.2). Student’s t-test was performed with parameter S0=0.1, FDR=0.05, number of randomizations=2500 in Perseus. Kinase-substrate datasets were obtained from PhosphoSitePlus and SIGNOR 3.0. Protein networks and pathway enrichments were analyzed using STRING [13].

Kinase activities were analyzed using the RoKAI app [14] with minimum substrate of 5 and minimum absolute Z-score of 2. The Kinase Library [15] was used for generation of the kinome tree. Phosphorylation motif enrichment analysis was conducted using iceLogo [16]. Analysis was performed based on precompiled Swiss-Prot composition with fold change scoring system and *p*-value of 0.05. Bar graphs and box plots were generated using GraphPad Prism 10.

## Results

### Differential proteomic and phosphoproteomic profiles in breast cancer cell lines

To achieve an integrative understanding of GC responses at protein and phosphoprotein level, three breast cancer cell lines were subjected to proteomic and phosphoproteomic analyses (Table 1). MCF7 and MDA-MB-231 cells were selected as two of the most widely used cell lines in breast cancer metastasis research [17], and MDA-MB-436 represents the protein dynamics associated with *BRCA1* mutation.

**Table 1.** Molecular information of breast cancer cell lines used in this study. TNBC (triple-negative breast cancer), WT (wild type), MT (mutant), ER (estrogen receptor), PR (progesterone receptor), HER2 (human epidermal growth factor receptor 2).

| Cell line | BRCA status | Type | TNBC |
| --- | --- | --- | --- |
| MCF7 | WT | Luminal A | No<br>(ER<sup>+</sup>/PR<sup>+</sup>/HER2<sup>-</sup>) |
| MDA-MB-231 | WT | Basal B | Yes |
| MDA-MB-436 | BRCA1 MT | Basal B | Yes |

State-of-the-art liquid chromatography tandem mass spectrometry (LC-MS/MS) using narrow-window data-independent acquisition (nDIA) mode [18] was employed to analyze the proteome and phosphoproteome changes in MCF7, MDA-MB-231, and MDA-MD-436 cells following 24-hour treatment of 0.1% DMSO or 1 µM Dex (Fig. 1a). The three cell lines exhibited clear separation by principal component analysis (PCA) of label-free MS-based protein and phosphosite intensities, respectively (Fig. 1b,c). Pairwise Pearson correlation coefficients were high within cell lines and as expected more variable for phosphoproteome than for proteome (Fig. 1d,e). Together, these results highlight global differences in protein abundance and greater divergence in phosphorylation-dependent signaling across cell lines.

Glucocorticoid receptor (GR) is known to be hyper-phosphorylated upon ligand binding, and its transcriptional activity is terminated by proteasomal degradation [19]. Pan-protein level of GR was significantly downregulated in all three cell lines after Dex treatment, however, phosphorylation of GR at Ser-267 and Thr-8 was less affected (Fig. 1f-h). Additionally, the basal level of total GR corresponded to Dex-mediated effect, as indicated by the counts of significantly regulated protein groups and phosphosites. MCF7 cells presented comparably low GR level compared to two other cell lines, resulting in substantially lower number of significantly regulated proteins (Fig. 1i-k). Based on this result, further analyses were performed for only MDA-MB-231 and MDA-MB-436 cells.

**Figure 1.**
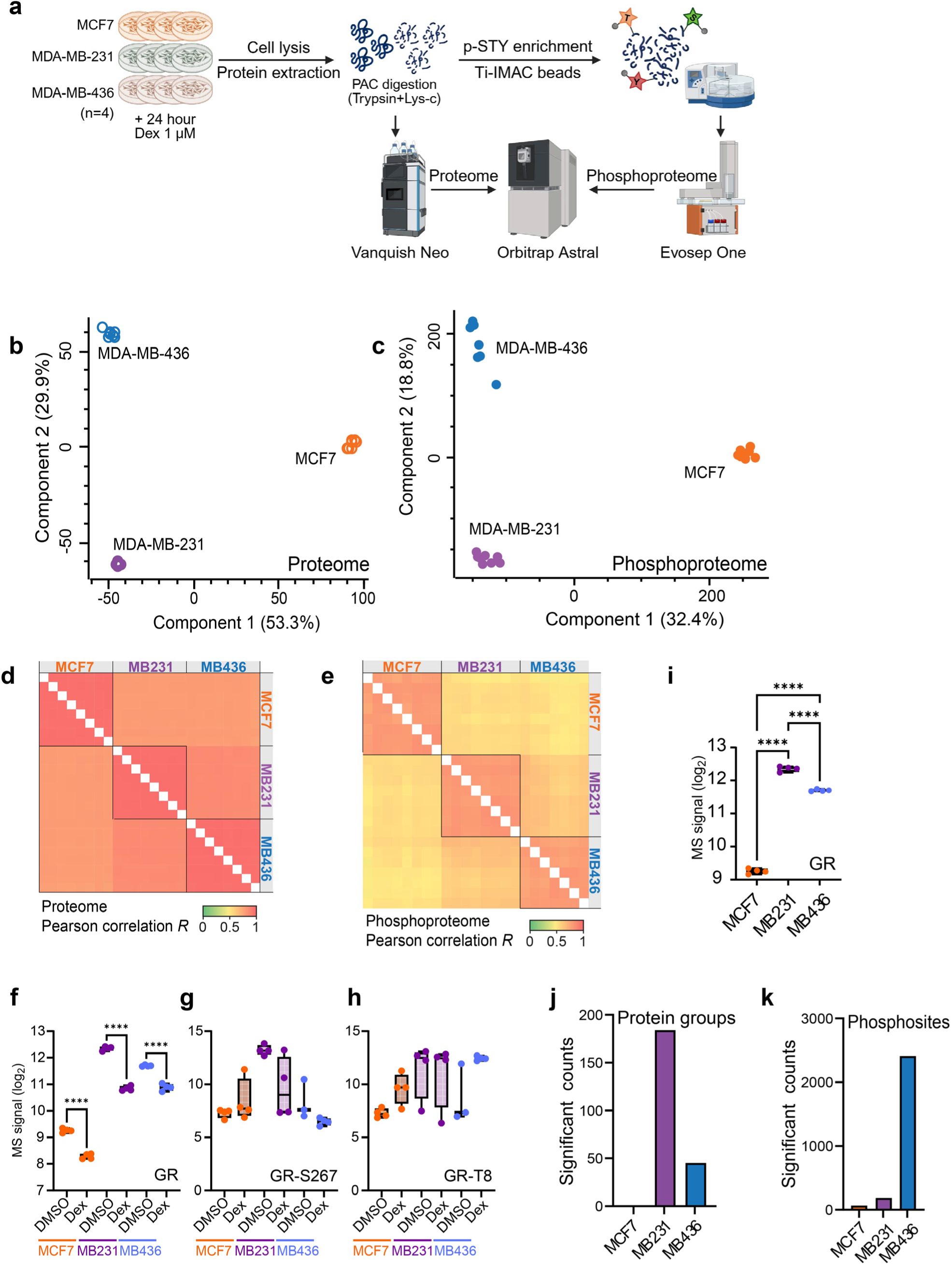
Distinct protein-level abundance in breast cancer cell lines a) A schematic workflow of quantitative mass spectrometry-based proteomic and phosphoproteomic analyses. Illustration was created using BioRender. b-c) PCA plot of proteome (b) and phosphoproteome (c) after 0.1% DMSO or 1 µM Dex treatment. d-e) Pearson correlation coefficient across all samples in proteome (d) and phosphoproteome (e). i) Box plot for individual log_2_ transformed label-free quantification of GR in breast cancer cell lines. f-h) Box plots for individual log_2_ transformed label-free quantification comparing DMSO and Dex treatment of GR (f), Ser-267 phosphorylation (g), and Thr-8 phosphorylation (h). Ordinary one-way ANOVA was used for statistical analysis. j-k) Bar plots showing counts of significantly regulated protein groups (j) and phosphosites (k).

### Dex treatment regulates proteins associated with metastasis in MDA-MB-231 cells

GR is a TF that can affect the activity of other TFs [20, 21]. Analyzing the proteome abundance changes using the TFLink database [22], we observed transcriptional regulatory interactions in the triple-negative MDA-MB-231 and MDA-MB-436 cells following Dex treatment. Among significantly up- or down-regulated protein groups in the proteome, 21 proteins were identified as TFs, including GR, SOX9, CEBPD, and PER1 that were shared in both cell lines (Fig. 2a). Notably, 13 of these TFs significantly modulated the abundance of 172 proteins identified as their transcriptional targets by TFLink in MDA-MB-231 cells (Fig. 2b). Dex-induced GR transcriptional activity was supported by pronounced fold-change of FKBP5, a well-established GR transcriptional target gene. Additionally, GR regulated the largest number of transcriptional targets among all TFs (Fig. 2c).

Next, functional pathway enrichment analysis was performed with the 172 proteins that were significantly regulated transcriptional targets of the 13 GC-responsive TFs in MDA-MB-231 cells. Top five pathways were selected based on signal using the STRING database [13] (Fig. 2d). WikiPathways revealed that these substrates were associated with GR pathway (WP2880) and nuclear receptors meta-pathway (WP2882). Additional enriched pathways included Gene Ontology (GO) biological processes such as regulation of cell adhesion (GO:0030155), positive regulation of cardiac EMT (GO:0062043), and regulation of cell motility (GO:2000145). Collectively, these findings suggest that Dex-induced GR alteration influences the activity of multiple TFs targeting GR-associated proteins in MDA-MB-231 cell line, which are also involved in breast cancer cell metastasis.

**Figure 2.**
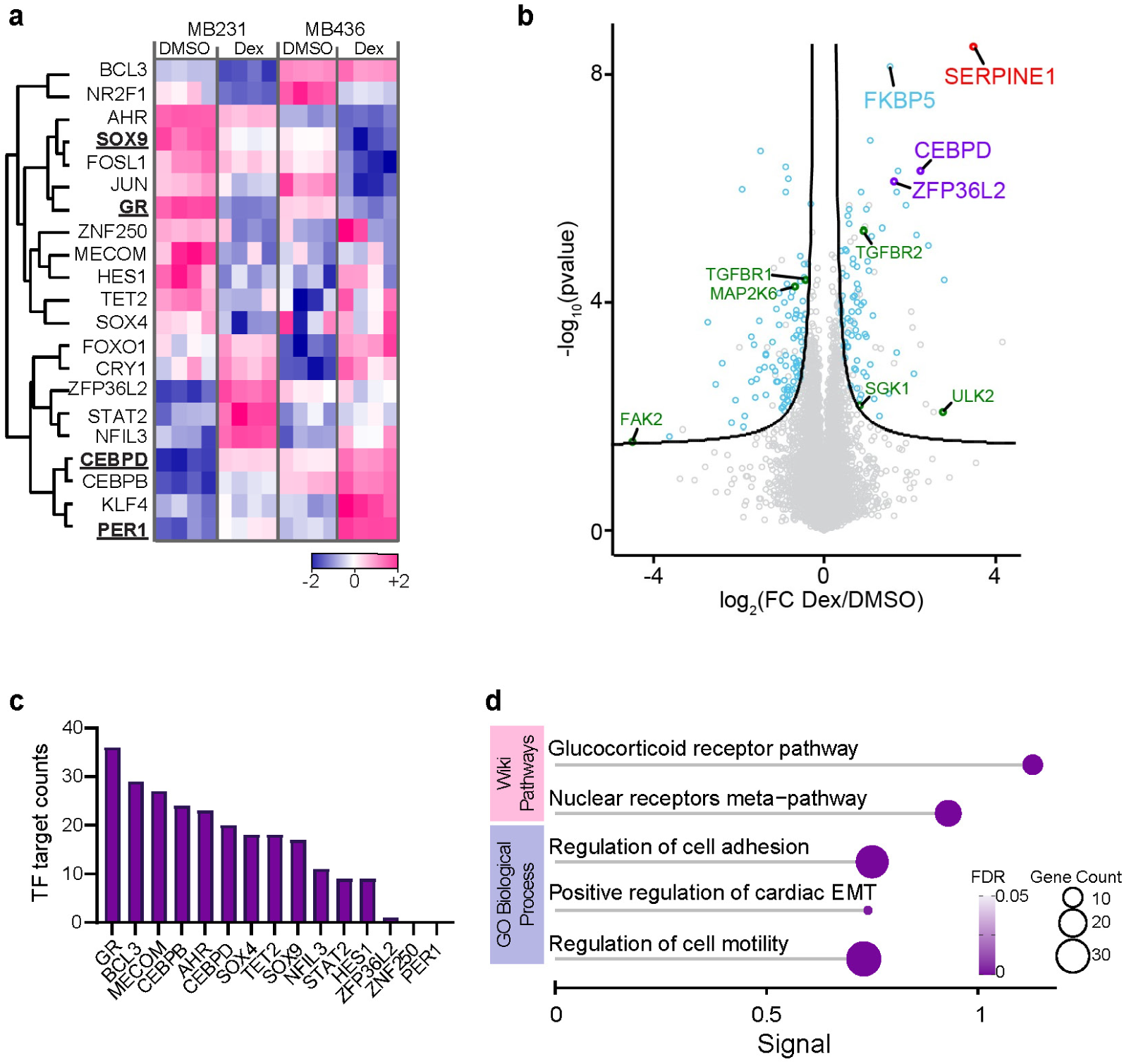
Proteomic changes in MDA-MB-231 cells after Dex treatment. a) Heatmap showing significantly Dex-regulated TFs in MDA-MB-231 and MDA-MB-436 cells. SOX9, GR, CEBPD, and PER1 are underlined to indicate overlapping regulation in both cell lines. b) Volcano plot of Dex-treated MDA-MB-231 proteome. Protein groups are colored as follows: transcriptional targets of significantly regulated TFs (blue), significantly phosphorylated TFs (purple), and transcriptional targets that are kinases (green). FC: fold-change. c) Bar plot showing the number of transcriptional targets per TF in MDA-MB-231 cells. d) Functional pathway enrichment analysis of 172 significantly regulated transcriptional targets in MDA-MB-231 cells.

### CK2 activation in MDA-MB-231 cells following Dex treatment

Among the 172 GC-regulated transcriptional targets in MDA-MB-231 cells, six were protein kinases of which ULK2, SGK1 and TGFBR2 were up-regulated whereas TGFBR1, MAP2K6, and PTK2B were down-regulated (Fig. 2b). Based on the kinase-substrate relationships acquired from PhosphoSitePlus (www.phosphosite.org) and SIGNOR 3.0 [23], none of their substrate proteins showed significant phosphorylation changes at known amino acid residues following Dex treatment.

Next, kinase activity analysis was performed employing the RoKAI algorithm [14]. The CK2 kinases were predicted to be significantly activated based on Dex-induced phosphorylation modifications in MDA-MB-231 cells (Fig. 3a). TTI1 and TOP2A were identified as direct phosphorylation substrates of CK2 kinases (Fig. 3b). TTI1 is not yet investigated in breast cancer, but has been reported to promote metastasis in non-small-cell lung cancer [24] and colorectal cancer [25]. On the other hand, TOP2A is a potential biomarker of breast cancer that correlates with poor prognosis [26–28]. Together, Dex-treated MDA-MB-231 cells reveal phosphorylation regulations that are associated with breast cancer metastasis.

**Figure 3.**
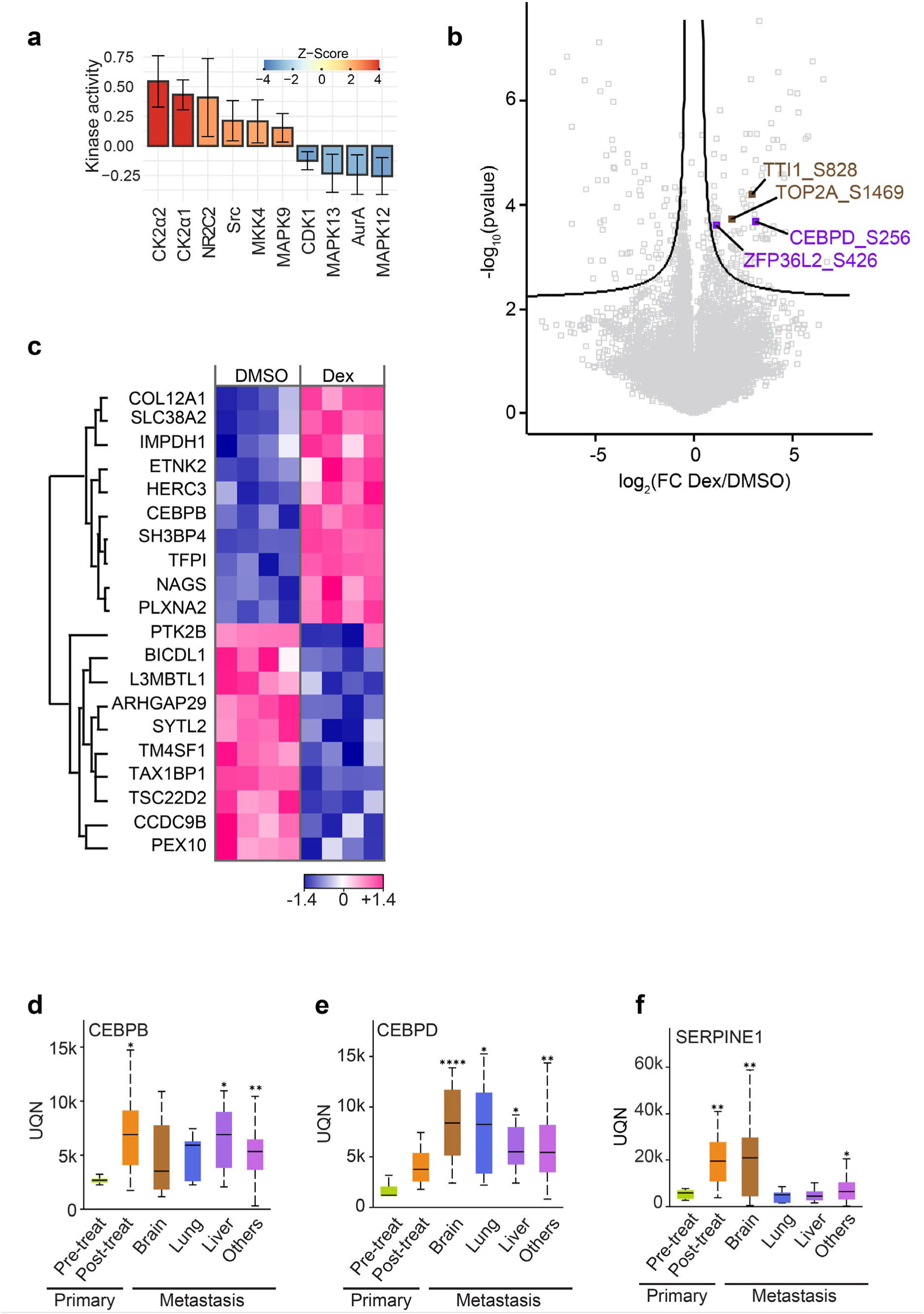
Phosphoproteomic changes in MDA-MB-231 cells after Dex treatment. a) Kinase activity analysis using the RoKAI app for MDA-MB-231 phosphoproteome after Dex treatment. b) Volcano plot of Dex-treated MDA-MB-231 phosphoproteome. Phosphosites are colored as follows: significantly phosphorylated TFs (purple) and CK2 substrates (brown). FC: fold-change. c) Heatmap showing differentially regulated CEBPD transcriptional targets after Dex treatment in MDA-MB-231 cells. d-f) Box plots showing CEBPB (d), CEBPD (e), and SERPINE1 (f) mRNA expressions in primary and metastasis breast cancer tumors. y-axis represents normalized RNA-seq samples fixed to upper quartile (UQN). Others include 16 sites that are not brain, lung, or liver. Post-treat refers to participants who received neoadjuvant therapy before primary tumor collection.

### Phosphorylation-mediated TF regulations in MDA-MB-231 cells

Since phosphorylation is an essential post-translational modification (PTM) for regulating the activity of many TFs [29], we examined changes in the identified 13 TFs at phosphorylation level and found that phosphorylation of CEBPD and ZFP36L2 increased at Ser-256 and Ser-426, respectively (Fig. 3b).

CCAAT-enhancer-binding proteins (C/EBPs) are a family of TFs consisting of six members, from alpha (α) to zeta (ζ). Although CEBPD is not a widely studied member of the family, elevated CEBPD is associated with chemotherapy resistance [30], cell proliferation [31], and stem cell-like phenotypes [32] of breast cancer. 20 protein groups were identified as transcriptional target genes of CEBPD in MDA-MB-231 proteome, including CEBPB that was significantly increased by Dex treatment (Fig. 3c).

Clinical relevance of CEBPB and CEBPD in breast cancer metastasis was evaluated using the AURORA US Metastatic Project dataset [33] available on MammOnc-DB [34]. Consistent with our hypothesis that GC treatment reprograms MDA-MB-231 proteome to promote metastasis, CEBPB and CEBPD mRNA expression levels were significantly higher in metastatic tumors than in primary tumors (Fig. 3d,e). Moreover, CEBPB expression correlated with lower 5-year survival probability in breast cancer patients from the SCAN-B project [35] (Supplementary Fig. 1a).

ZFP36L2 was another TF that exhibited significant phosphorylation-level upregulation in MDA-MB-231 cells. ZFP36L2 has been implicated in tumorigenesis of many cancers [36] and is associated with bone metastasis in breast cancer [37]. According to the TFLink database, ZFP36L2 regulates SERPINE1 transcription. Interestingly, SERPINE1 was the most substantially upregulated protein in MDA-MB-231 proteome after Dex treatment (Fig. 2b). The AURORA dataset presented elevated mRNA expression level of SERPINE1 in metastasis tumors in brain and other organs (Fig. 3f). High expression of SERPINE was also associated with reduced survival in breast cancer patients (Supplementary Fig. 1b).

**Supplementary Figure S1.**
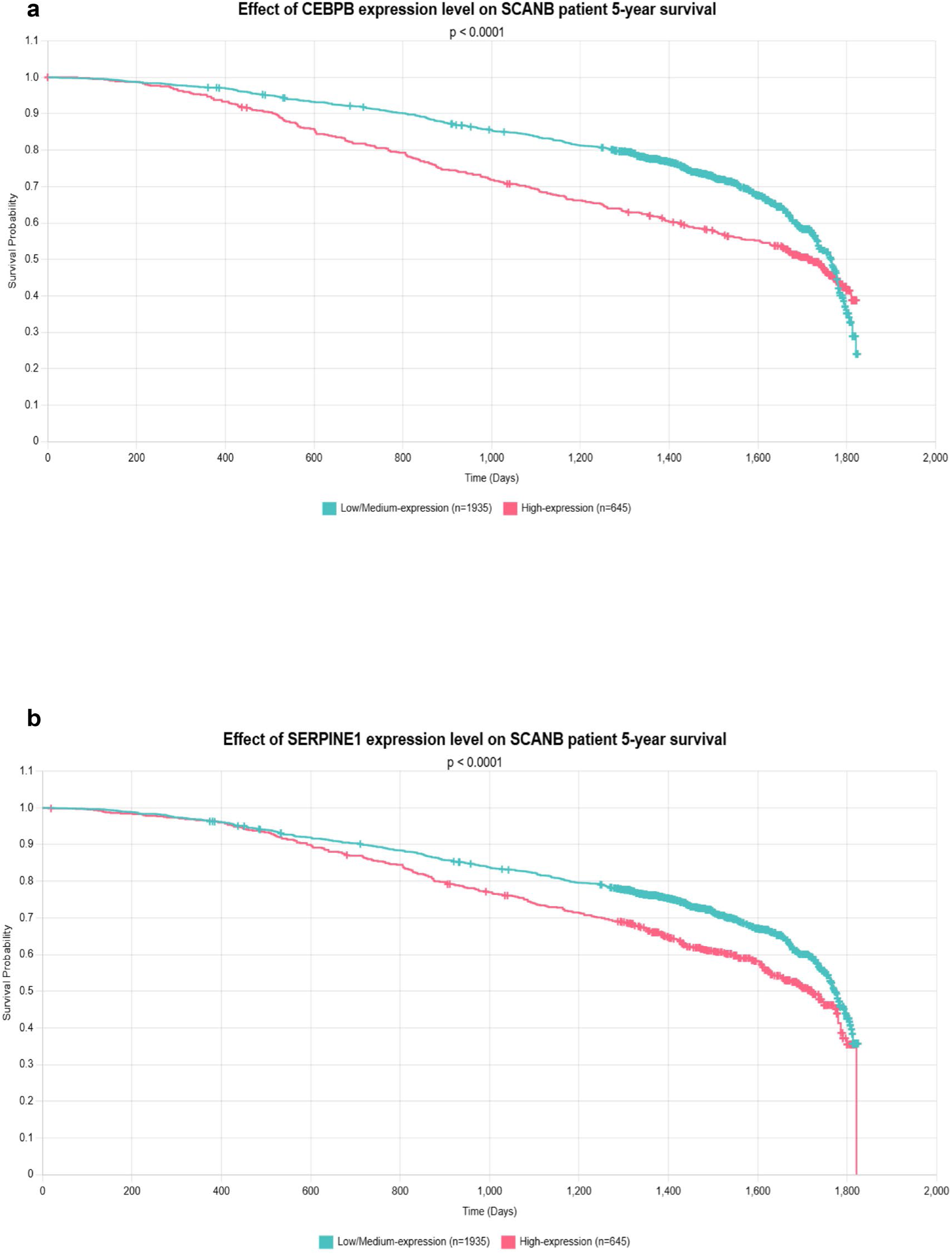
Survival plot of SCAN-B patients. a-b) Effect of CEBPB (a) and SERPINE1 (b) expression in 5-year survival of breast cancer patients from SCAN-B project.

### Dex-induced GR transcriptional activity in MDA-MB-436 cells

To assess the differences driven by *BRCA1* mutation in TNBC cells, the effect of Dex treatment was investigated in MDA-MB-436 cells. The same computational approach based on the TFLink database was implemented to identify 37 transcriptional target genes significantly regulated by 10 TFs in the proteome profile. Interestingly, GR was not the top-most regulating TF, but prominent increase of FKBP5 abundance confirmed Dex-mediated effect in MDA-MB-436 cells (Fig. 4a). Similar to the result of MDA-MB-231 cells, functional pathway enrichment analysis by STRING revealed that these transcriptional targets were enriched in GC-associated signaling pathways including GR pathway, nuclear receptors meta-pathway, and negative regulation of GR signaling pathway (GO:2000323). Interestingly, they were also involved in melatonin metabolism and effects (WP3298) and response to insulin (GO:0032868) (Fig. 4b).

**Figure 4.**
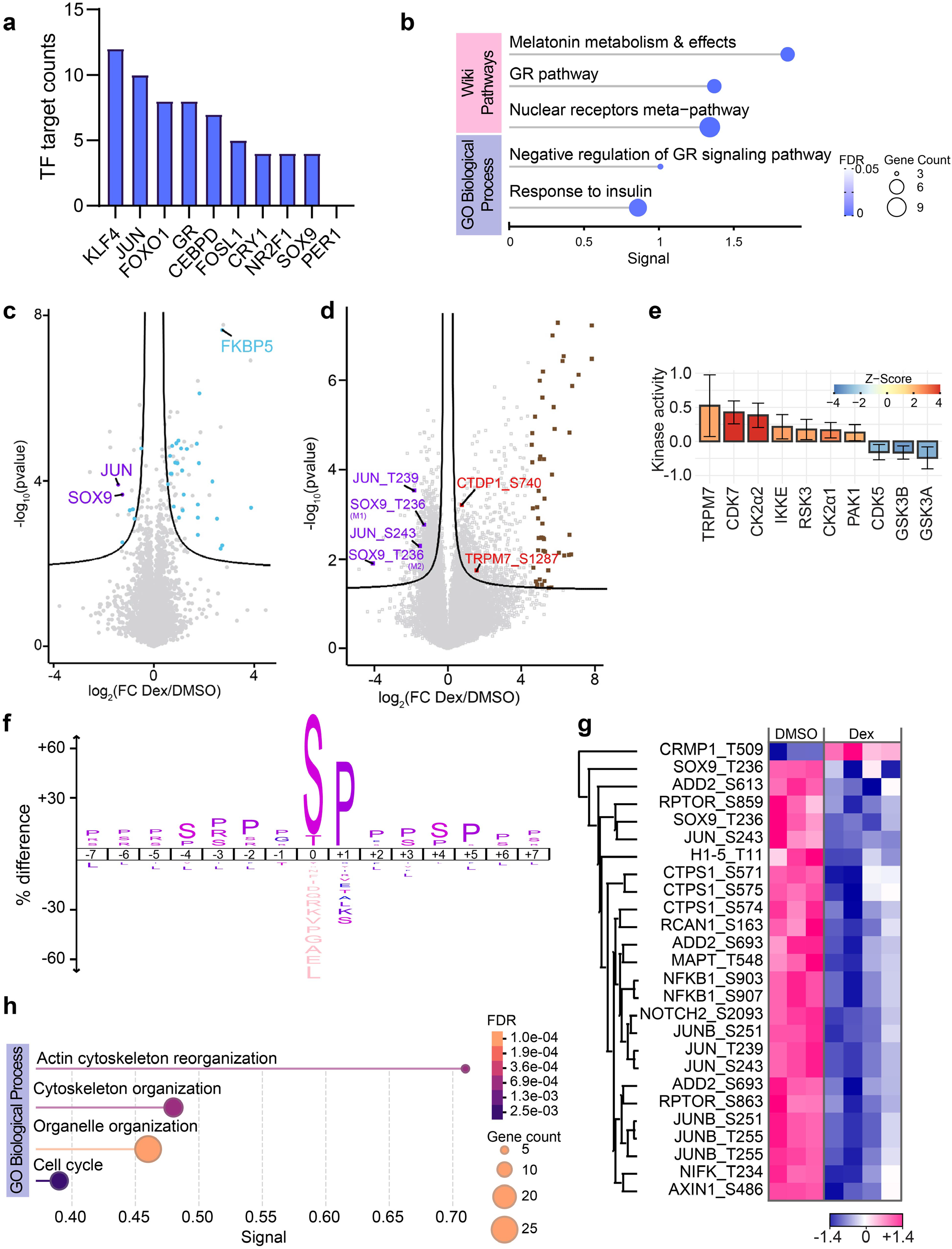
Proteomic and phosphoproteomic changes in MDA-MB-436 cells after Dex treatment. a) Bar plot showing the number of transcriptional targets per TF in MDA-MB-436 cells. b) Functional pathway enrichment analysis of 37 significantly regulated transcriptional targets in MDA-MB-436 cells. c-d) Volcano plots of Dex-treated MDA-MB-231 proteome (c) and phosphoproteome (d). Protein groups or phosphosites are colored as follows: transcriptional targets of significantly regulated TFs (blue), top 50 upregulated phosphosites (brown), and significantly phosphorylated TFs (purple). FC: fold-change. e) Kinase activity analysis using the RoKAI app for MDA-MB-436 phosphoproteome after Dex treatment. f) Motif enrichment analysis of significantly downregulated phosphosites using iceLogo. g) Heatmap for significantly regulated direct substrates of GSK3A and GSK3B. h) Functional pathway enrichment analysis of top 50 upregulated phosphosites.

### GSK3A/B activity suppression by Dex in MDA-MB-436 cells

Next, we performed an integrative analysis combining the proteome and phosphoproteome data from MDA-MB-436 cells. None of the 37 transcriptional targets with significant abundance changes in the proteome were identified as kinases. Among significantly phosphorylation-regulated TFs, phosphorylation level of SOX9 Thr-236, JUN Thr-239, and JUN Ser-243 were suppressed by Dex (Fig. 4d). All of these three phosphosites are direct substrates of GSK3A and GSK3B, whose activity was predicted to be suppressed by the RoKAI algorithm (Fig. 4e). Conversely, the total and phosphorylated level of GSK3 proteins were not affected by Dex (Supplementary Fig. 2a,b), indicating post-translational regulation of GSK3 activity after GC-treatment. The flanking region of all 1709 significantly downregulated phosphosites were enriched for the pS-xxx-S motif, which is a conserved recognition motif of GSK3 (Fig. 4f) [38]. GSK3 substrates were significantly downregulated except for CRMP1 in the phosphoproteome, further supporting inactivation of GSK3 kinases (Fig. 4g).

**Supplementary Figure S2.**
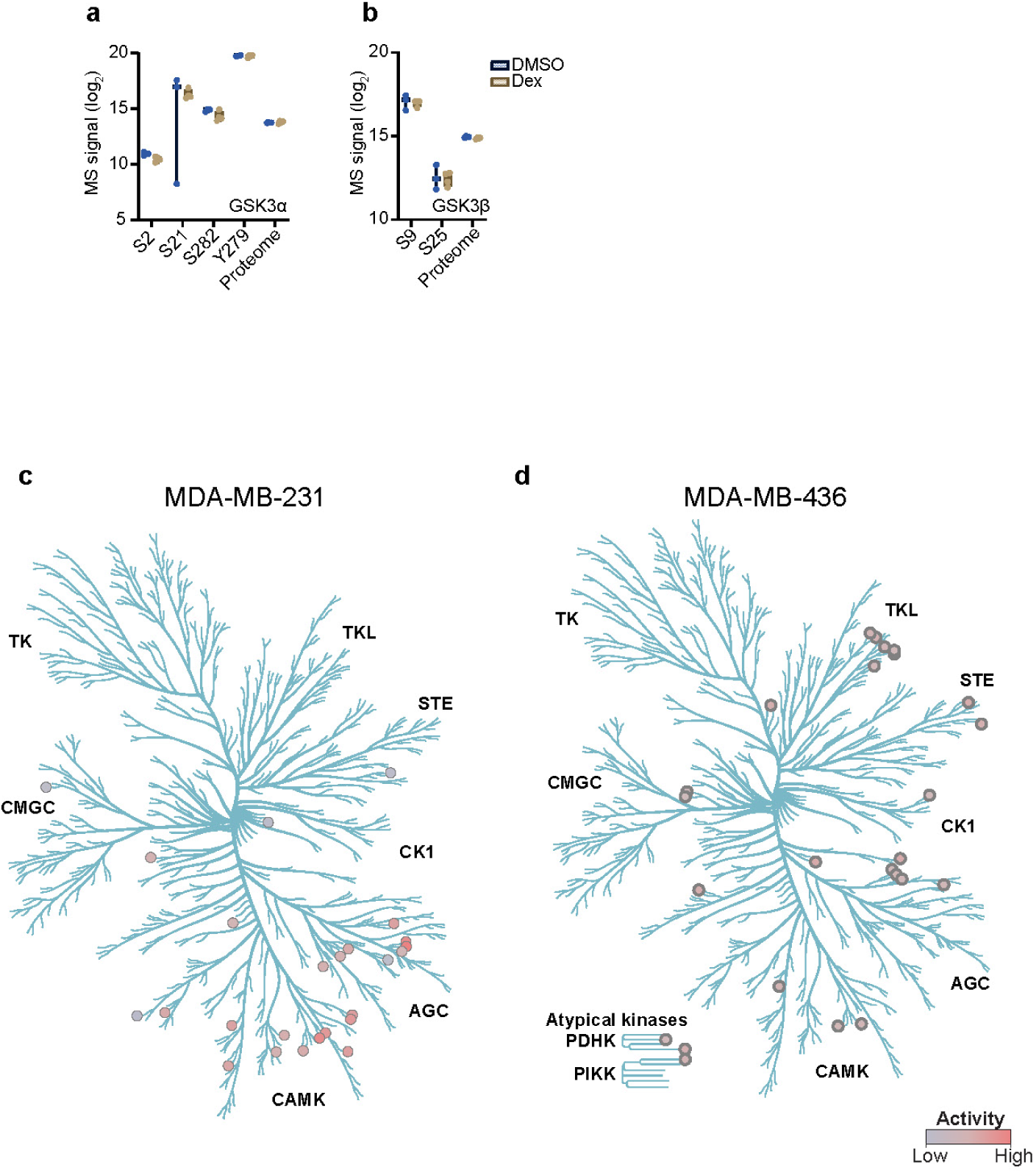
Dex-induced phosphorylation changes in MDA-MB-231 and MDA-MB-436 cells. a-b) Box plots for individual log_2_ transformed label-free quantification of GSK3 total protein and phosphorylation in breast cancer cell lines. c-d) Kinome tree of Dex-treated breast cancer cell lines.

### Dex treatment promotes phosphorylation of metastasis-associated proteins in MDA-MB-436 cells

We then examined the upregulatory effect of Dex treatment in MDA-MB-436 cells. Among the kinases predicted to be activated by RoKAI algorithm, TRPM7 autophosphorylated at Ser-1287 while CK2 induced phosphorylation of CTDP1 at Ser-740 (Fig. 4e). TRPM7 utilizes its kinase domain to regulate Src and MAPK signaling, ultimately affecting breast cancer cell metastasis [39–41]. CTDP1, as the only phosphatase identified in the human proteome with *BRCA1* C-terminal domain, promotes breast cancer cell survival through efficient homologous recombination repair [42].

To obtain a more comprehensive understanding of phosphorylation-mediated stimulation by Dex, 48 protein groups from the top 50 upregulated phosphosites were analyzed by STRING. These proteins were involved in GO biological processes related to actin cytoskeleton (GO:0007010, GO:0031532) and organelle organization (GO:0006996) (Fig. 4h), suggesting that Dex treatment remodels global protein abundance relevant to cell metastasis regardless of *BRCA1* mutation.

## Discussion

Breast cancer is a prevailing cause of cancer-related death in females worldwide. Its treatment strategies include surgery, chemotherapy, targeted therapy, and radiotherapy [2]. To prevent the known side effects of these treatments such as vomiting and nausea, systemic GCs are often prescribed in combination with conventional drugs [8]. In this study, we acquired proteome and phosphoproteome of breast cancer cells after Dex treatments to better understand the regulatory mechanisms of GCs. To date, this is the first study that compares protein-level signatures of Dex responses in multiple breast cancer cell lines.

First, we confirmed the biological diversity of MCF7, MDA-MB-231, and MDA-MB-436 cell lines in both proteome and phosphoproteome profiles. The number of significantly regulated protein groups and phosphosites following Dex treatment depended on the basal level of GR expression. Significantly regulated protein and phosphosite counts were comparably negligible with zero and 67, respectively, in MCF7 cells. This is likely explained by the substantially lower protein abundance level of GR in MCF7 cells compared to MDA-MB-231 and MDA-MB-436 cells.

MDA-MB-231 and MDA-MB-436 cells displayed distinct proteome and phosphoproteome profiles after Dex treatment. It remains unclear why MDA-MB-231 cells exhibited extensive abundance changes at the proteome level but comparatively fewer phosphoproteome alterations relative to MDA-MB-436 cells. This divergence was also reflected in kinase activity analysis, where MDA-MB-436 cells showed broader global kinase modulation. On the other hand, the affected kinases in MDA-MB-231 cells were predominantly members of the AGC and CAMK families (Supplementary Fig. 2c,d). This may be due to differences in *BRCA1* mutation, but additional validation is required to confirm cell line-specific responses.

Ensuing, we demonstrate that Dex treatment modulated the activity of multiple TFs in MDA-MB-231 and MDA-MB-436 cell lines. As expected, transcriptional activity of GR was promoted after the treatment, indicated by upregulation of FKBP5 protein levels and functional pathway enrichment in GR-associated signaling. Hyper-phosphorylation of GR, however, was not detected in the phosphoproteome. This may be due to missing phosphorylation signals for its functionally known residues, Ser-203, Ser-211, and Ser-226 [43]. Further studies are necessary to clarify whether Ser-267 and Thr-8 residues of GR relate to its transcriptional activity.

Here, we suggest several noncanonical TFs that are regulated by Dex treatment. In MDA-MB-231 cells, phosphorylation of CEBPD and ZFP36L2 significantly increased, and their transcriptional target genes were overexpressed in distant breast cancer tumors. Transcriptional activity of CEBPD can be enhanced by phosphorylation at Ser-167 by GSK3B [44] and at Ser-191 by PKA [45]. Consistent with this, we observed upregulated protein abundance levels of 10 transcriptional target genes of CEBPD including CEBPB, which is a well-recognized factor that promotes breast cancer metastasis [46–49]. However, further investigation is necessary to determine whether phosphorylation of CEBPD at Ser-256 residue impacts its transcriptional activity, as protein abundance of 10 transcriptional targets were significantly downregulated.

In MDA-MB-231 cells, ZFP36L2 was another TF that was phospho-upregulated after Dex treatment. ZFP36L2 can be phosphorylated by LPS stimulation [50] or ERK [51], thereby promoting mRNA stabilization. Again, additional studies are required to unveil the role of Ser-426 phosphorylation at ZFP36L2. SERPINE1, a transcriptional target of ZFP36L2 according to the TFLink database, outstandingly increased at protein level following Dex treatment. Recent reports underscore the function of SERPINE1 in driving aggressiveness, chemotherapy resistance, and metastasis of breast cancer [52–54]. Collectively, these findings support our hypothesis that Dex treatment promotes metastasis in MDA-MB-231 cell line.

Our data suggest that GSK3 catalytic kinase activity was greatly suppressed in Dex-treated MDA-MB-436 cells, thereby downregulating phosphorylation of JUN and SOX9 among significantly regulated TFs. GSK3 is a potential target of metastatic breast cancer and GSK3B inhibitors can suppress mesenchymal and stem cell-like properties in TNBC [55, 56]. In this sense, systemic GC treatment may be beneficial. However, proteins that displayed increased phosphorylation were enriched in signaling pathways associated with cytoskeletal organization, implying that cells can be more migratory. *In vitro* cell-based assays can be performed to validate cell migration and invasion in breast cancer for additional analyses.

In this study, we provide a useful resource for the breast cancer research community by comparing Dex responses in three breast cancer cell lines with distinct molecular features. Our data suggest that Dex treatment regulates proteins associated with breast cancer metastasis. However, it is crucial to note that this study focuses on protein-level modulations and further molecular studies are needed to confirm the underlying biological mechanisms.

## Acknowledgements

Work at the Novo Nordisk Foundation Center for Protein Research (NNF CPR) is funded in part by a donation from the Novo Nordisk Foundation (NNF14CC0001 and NNF24SA0098829). This work has been funded by Independent Research Fund Denmark | Natural Sciences under the grant agreement no. 2032-00311B. This project was supported by a center-of-excellence grant from the Danish National Research Foundation to Copenhagen Center for Glycocalyx Research (DNRF196). The project was also supported by a generous grant from the Danish Agency of Higher Education and Science to establish the PLATO research infrastructure: Danish National Mass Spectrometry Platform for Proteomics and Biomolecular Imaging (grant no. 5229-00012B). The authors would like to thank NNF CPR Mass Spectrometry Platform for MS instrument and technical assistance. The authors have declared no competing interest.

## Author contributions

HC designed the study, acquired and analyzed data, drafted the original manuscript, and generated the figures. JVO provided resources and supervision. HC and JVO read and edited the final manuscript.

## Data availability

All mass spectrometry raw files and Spectronaut searches are available on the ProteomeXchange Consortium via the PRIDE repository with the dataset identifier PXD070780.

